# ZW sex-chromosome evolution and contagious parthenogenesis in Artemia brine shrimp

**DOI:** 10.1101/2022.04.27.489735

**Authors:** Marwan Elkrewi, Uladzislava Khauratovich, Melissa A. Toups, Vincent Kiplangat Bett, Andrea Mrnjavac, Ariana Macon, Christelle Fraisse, Luca Sax, Ann Kathrin Huylmans, Francisco Hontoria, Beatriz Vicoso

## Abstract

Eurasian brine shrimp (genus *Artemia*) have closely related sexual and asexual lineages of parthenogenetic females, which produce rare males at low frequencies. Although they are known to have ZW chromosomes, these are not well characterized, and it is unclear whether they are shared across the clade. Furthermore, the underlying genetic architecture of the transmission of asexuality, which can occur when rare males mate with closely related sexual females, is not well understood. We produced a chromosome-level assembly for the Eurasian species *A. sinica* and characterized in detail the pair of sex chromosomes of this species. We combined this with short-read genomic data for the sexual species *A. sp. Kazakhstan* and several lineages of *A. parthenogenetica,* allowing us to perform a first in-depth characterization of sex-chromosome evolution across the genus. We identified a small differentiated region of the ZW pair that is shared by all sexual and asexual lineages, supporting the shared ancestry of the sex chromosomes. We also inferred that recombination suppression has spread to larger sections of the chromosome independently in the American and Eurasian lineages. Finally, we took advantage of a rare male, which we backcrossed to sexual females, to explore the genetic basis of asexuality. Our results suggest that parthenogenesis may be partly controlled by a locus on the Z chromosome, highlighting the interplay between sex determination and asexuality.

## Introduction

The diversity of reproductive and sex-determining systems has long puzzled evolutionary biologists (Bachtrog *et al*. 2014; Pennell *et al*. 2018; Picard *et al*. 2021). When separate sexes are present, the development of males and females can be controlled by environmental factors or through the presence of sex-determining loci (Beukeboom and Perrin 2014; Bachtrog *et al*. 2014). These sex determining loci are typically carried by specialized “sex chromosomes”, such as the X and Y chromosomes of mammals. Sex chromosomes initially arise from standard pairs of autosomes, but can progressively stop recombining over much of their length, ultimately resulting in genetic and morphological differentiation (Charlesworth *et al*. 2005; Wright *et al*. 2016). Each segment of the sex chromosome pair that stopped recombining at a given timepoint is referred to as a “stratum”, and strata of different ages are often found on the same pair of sex chromosomes (Lahn and Page 1999; Handley *et al*. 2004). The Y chromosome stops recombining altogether after XY recombination suppression and eventually degenerates, i.e. it accumulates deleterious mutations and can lose many or even all of its genes (Bachtrog 2013). This gene loss leads to dosage deficits in males, since many X-linked genes become single-copy. Mechanisms of dosage compensation often target the X-chromosome and regulate its expression, thereby reestablishing optimal dosage balance of genes across the genome (Charlesworth 1978; Straub and Becker 2007; Vicoso and Bachtrog 2009; Disteche 2016). Much of our understanding of these processes has come from studying the ancient XY systems of traditional model organisms such as mice and fruit flies. Despite the recent characterization of young sex chromosomes in many nonmodel species (Charlesworth 2019), many questions remain about the earlier stages of sex-chromosome divergence, such as what molecular mechanisms and selective pressures drive the initial loss of recombination between sex chromosomes (Ponnikas *et al*. 2018). Similarly, female-heterogametic species (i.e. females are ZW, males are ZZ) have remained relatively understudied, as they are not found in any of the main model organisms. While parallels exist between the evolution of XY and ZW pairs, such as the progressive loss of recombination and subsequent degradation of the Y/W-chromosomes (Ellegren 2011; Vicoso *et al*. 2013; Zhou *et al*. 2014; Picard *et al*. 2018; Sigeman *et al*. 2021), some aspects of their evolution seem to differ. In particular, dosage compensation of Z-chromosomes is often limited to a few dosage-sensitive genes (i.e. it works gene-by-gene, as opposed to the chromosome-wide mechanisms found in many XY species, (Mank 2013; Rovatsos and Kratochvíl 2021)). These discrepancies may have to do with systematic differences in selection and mutation between males and females (Vicoso and Bachtrog 2009; Ellegren 2011; Mullon *et al*. 2015), or may simply be a coincidence due to the few ZW systems characterized in detail at the molecular level (Rovatsos and Kratochvíl 2021).

Although the prevalence of sexual reproduction suggests that it offers long-term advantages, asexual lineages are found in many clades and successfully inhabit a variety of ecological niches (Toman and Flegr 2018). Transitions from sexual to asexual reproduction are frequent (Neiman et al., 2014), and can involve a diversity of mechanisms that disrupt meiosis, such as novel mutations, hybridization of closely related lineages, and polyploidization (Neiman *et al*. 2014). Asexuality can evolve from any ancestral sex-determining system, including in species with differentiated sex chromosomes, (e.g. Schwander and Crespi 2009; Jaquiéry *et al*. 2014; Mignerot *et al*. 2019), and understanding the mechanisms underlying these transitions has been a key goal of the field.

In many asexual lineages, males are occasionally produced, and can fertilize closely related sexual females, which then give rise to new asexual lineages (“contagious parthenogenesis”). These crosses have facilitated the use of traditional genetic approaches for understanding the genetic architecture of asexuality (Jaquiéry *et al*. 2014). Transitions from sexual to asexual reproduction have primarily been studied in animal species where both sexual reproduction and parthenogenesis were ancestrally part of the life cycle, either in the form of cyclical parthenogenesis or haploidiploidy (Neiman *et al*. 2014). In this case, the loss of sexual reproduction and consequent obligatory parthenogenesis is often controlled by one or only a few loci (Lynch *et al*. 2008; Sandrock and Vorburger 2011; Eads *et al*. 2012; Jaquiéry *et al*. 2014; Aumer *et al*. 2017; Yagound *et al*. 2020). In the pea aphid, the locus controlling asexuality is found on the X-chromosome (Jaquiéry *et al*. 2014), and a locus of large effect on parthenogenesis was also found on the UV sex chromosome pair of brown algae *Ectocarpus* (Mignerot *et al*. 2019), raising interesting questions about the interplay between the ancestral sex-determining system and contagious parthenogenesis. One direct link between the two phenomena is that when asexuals are derived from an ancestral XX/XY or haplodiploid sex-determination systems, rare males can be formed through the loss of an X-chromosome (Kampfraath *et al*. 2020) or through accidental production of haploid individuals during automixis (Sandrock and Vorburger 2011). Less is known about the creation of rare males when the ancestral sex-determination system was female-heterogamety. More generally, it is unclear if sex chromosomes are a prime spot for the location of genes regulating asexual reproduction, since very few transitions have been characterized in organisms with sex chromosomes.

Brine shrimp of the genus *Artemia* have both asexual and sexual species (Abatzopoulos 2018), as well as ZW sex chromosomes with putative ancient and recent strata (Bowen 1963; De Vos *et al*. 2013; Accioly *et al*. 2015; Huylmans *et al*. 2019), making them an ideal model for addressing many of these questions. While all new world species are sexual, the Eurasian clade consists of a few sexual species (including *A. sinica, A. sp. Kazakhstan* and *A. urmiana*) and of various asexual lineages (collectively refered to as *A. parthenogenetica*, and further referred to by their location of origin; Van Stappen 2002; Maccari *et al*. 2013b). Asexuals vary in ploidy, but only diploid lineages are considered here. Originally thought of as fairly ancient, these lineages turned out to have arisen recently through hybridization between asexual lineages and individuals from or closely related to *A. sp. Kazakhstan* (Baxevanis *et al*. 2006; Maccari *et al*. 2013b; Rode *et al*. 2021). In *Artemia*, such contagious parthenogenesis can occur through the production of rare males by asexual lineages, which can fertilize closely related sexual females (Maccari *et al*. 2013a; Abatzopoulos 2018). Furthermore, asexual females can mate with males of sexual species and produce a minority of offspring sexually (Boyer *et al*. 2021). The ZW pair of *Artemia* has been mostly studied in the American species *A. franciscana* (Bowen 1963; Parraguez *et al*. 2009; De Vos *et al*. 2013; Accioly *et al*. 2015). Both a small differentiated region and a non-recombining but largely undifferentiated region were detected, making it an interesting system to understand the first steps leading to ZW divergence (Huylmans *et al*. 2019). Gene expression in the differentiated region appears to be fully balanced between males and females, but there was limited power to detect changes due to the fragmented nature of the genome (Huylmans *et al*. 2019). Eurasian lineages also carry a ZW pair (Haag *et al*. 2017), but whether the same chromosome is used for sex determination accross the clade in not known. Because *A. parthenogenetica* reproduce through central fusion automixis (Nougué *et al*. 2015), a modified form of meiosis, which allows for loss of heterozygosity when recombination between chromosomes occurs, rare recombination events between the Z and W (which replace part of the W with its Z-linked homologous region) can lead to the creation of rare males (Nougué *et al*. 2015; Boyer *et al*. 2022). Finally, the genetic mechanisms behind asexuality, and whether the sex chromosomes play any further role in its evolution, have not yet been explored in detail.

Here, we develop several genomic resources for *Artemia* lineages, including the first chromosome-level assembly for the *Artemia* genus (*A. sinica*), as well as short-read genomic data for *A. sp. Kazakhstan* and several lineages of *A. parthenogenetica*. Using these data, we are able to provide the first in-depth characterization of sex-chromosome evolution across the genus, including identifying an ancient region shared with the American species *A. franciscana*. Finally, we find evidence that asexuality is likely partly controlled by a locus on the Z chromosome - a first in a ZW sex chromosome system.

## Results

### 1. The ZW pair is shared by American and Eurasian *Artemia*

Two genome assemblies and a high density linkage map are currently available for the American *A. franciscana* (Jo *et al*. 2021a; Han *et al*. 2021; De Vos *et al*. 2021), but resources for the Eurasian clade are more limited, with only an *A. sp. Kazakhstan* draft genome assembly recently described in Boyer et al. (2022). The median dS between the two clades is ∼0.2. We assembled a male genome of *A. sinica* using PacBio long reads (∼30x) and Hi-C Illumina reads (1.5*e12 reads), yielding 1213 scaffolds with an N50 of 67.19 Mb (Sup. Fig. 1) and a total length of 1.7Gb; 85% of the sequences get assigned to one of the 21 largest scaffolds (which corresponds to the expected number of chromosomes, Sainz-Escudero *et al*. 2021). The strong diagonal in the heatmap of the Hi-C contact matrix (Sup. Fig. 2) supports the high quality of our assembly, as does our BUSCO score of 91.8%. This chromosome-level assembly represents an improvement over existing *Artemia* genomes, which have N50 values of 27 to 112Kb, and BUSCO scores of 68.3% to 86.9% (Jo *et al*. 2021a; De Vos *et al*. 2021; Boyer *et al*.; Sup. Fig. 3).

Our earlier analysis of female and male genomic coverage in *A. franciscana* had uncovered a small region of reduced female coverage, consistent with full differentiation of the Z and W chromosomes (Huylmans *et al*. 2019). To investigate whether ZW differentiation was also present in *A. sinica*, we first estimated male and female coverage along each chromosome. Consistent with *A. franciscana*, only a small genomic region on chromosome 1 had decreased female/male coverage (Fig. 1A, Sup. Fig. 4 for all chromosomes), showing that chromosome 1 is the Z chromosome. To check for homology with the *A. franciscana* differentiated region, we mapped the scaffolds from the *A. franciscana* genome of (Jo *et al*. 2021a) to the new *A. sinica* assembly based on their shared gene content, and plotted the coverage values that we had previously estimated (Huylmans *et al*. 2019) based on the *A. sinica* coordinates. Fig. 1A shows that the two differentiated regions largely overlap, supporting the ancestry of the pair of sex chromosomes; we name this shared region stratum 0 (S0). In the *A. franciscana* linkage map (Han *et al*. 2021), LG6 was identified as the sex chromosome. To further verify the homology between the ZW pairs of the two species, we mapped the genetic markers used by Han et al. (2021) to our *A. sinica* assembly. As expected, the vast majority of LG6 markers for which we could infer a location mapped to our chromosome 1 (Sup. Fig. 5). We also produced an assembly based on female long PacBio reads, which contains a substantial amount of scaffolds with excessive female coverage, consistent with W-linkage (Sup. Fig. 6).

**Fig. 1:**
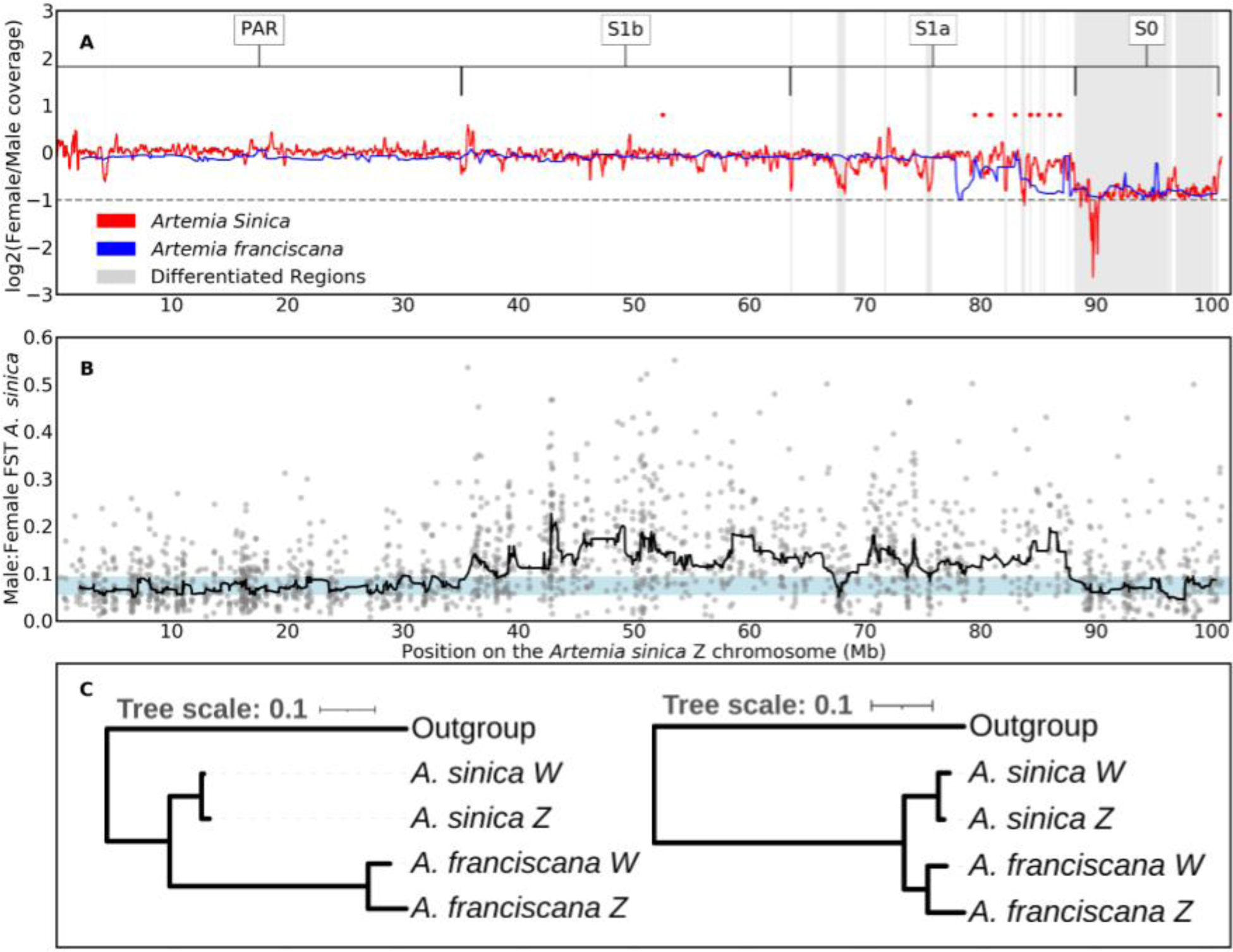
A shared sex-linked region on the ZW pair. A. Patterns of female/male coverage in *A. franciscana* and *A. sinica*. The differentiated regions are highlighted in gray, and the putative strata are defined above. The red dots are the locations of the W-candidates shared between *A. sinica* and *A. franciscana*. The horizontal gray line is at -1 to signify the regions with a twofold reduction in coverage in females compared to males. B. Male:female FST along the putative chromosome Z. The dots are FST calculated for bins of 1000 nucleotides, and the black line is the rolling median computed in sliding windows of 30 consecutive 1000 nucleotide bins. The light blue shaded area highlights the region between the 5^th^ and 95^th^-percentiles of the rolling median of FST for autosomes. C. Phylogenetic trees for two examples of the W-candidates shared between *A. sinica* and *A. franciscana* and their putative Z homologs. *Branchinecta lindahli* is used as the outgroup.

### 2. Convergent loss of ZW recombination

To identify parts of the sex chromosomes that no longer recombine, but are still similar enough that W-derived reads still map to the Z, we used previously published RNA-seq dataset for *A. sinica* (Huylmans *et al*. 2021), obtained from 10 males and 10 females, to estimate FST, a measure of genetic differentiation, between the two sexes. Genetic variants found exclusively on the W increase the level of female-male differentiation, and young non-recombining regions can be detected through their high male:female FST (Palmer *et al*. 2019; Vicoso 2019; Gammerdinger *et al*. 2020). Fig. 1B shows that a large region (∼52 Mb) has FST values systematically above the 95^th^-percentile of autosomes, consistent with recent loss of recombination in *A. sinica*. We call this region Stratum 1 (S1), but further divide it into S1a, which shows localized drops in female:male coverage (gray shaded regions in Fig. 1A), and S1b, for which no coverage differences are observed (Fig. 1A).

Given the substantial distance between the two lineages, we hypothesized that the loss of recombination in S1 had occurred specifically in the Eurasian clade. To verify this, we used a k-mer-based pipeline combining male and female DNA and RNA short reads (Elkrewi *et al*. 2021) to identify putative W-derived transcripts. This yielded 402 transcripts in *A. franciscana* and 319 in *A. sinica*. Of those that mapped to the genome, 182 out of 310 (59%) *A. sinica* transcripts and 168 out of 364 (46%) *A. franciscana* transcripts mapped to chromosome 1 (Z) of *A. sinica*, a higher proportion than the overall 7% of genes that map to this chromosome, confirming the validity of the approach (since we expect many W-linked genes to have a close homolog on the Z). Few of these candidate W genes mapped to the putative ancestral sex-linked region (16 in *A. sinica*, compared to 84 genes in the corresponding Z-linked region), consistent with substantial degeneration of this part of the W-chromosome. To find genes present on the W-chromosomes of both species, we selected reciprocal best hits between the two sets of W candidates. Surprisingly, 15 were found in both species, but mapped to the putative S1a region, suggesting that at least part of this region has also stopped recombining in *A. franciscana*. We made phylogenetic trees using each pair of homologous W-genes and their Z-linked homologs (as well as the closest homolog obtained from the transcriptome of the distantly related fairy shrimp *Branchinecta lindahli* (Schwentner *et al*. 2018), when one could be detected, to infer whether they were W-linked before the split of the two clades. ZW homologs clustered by species rather than by chromosome (Fig. 1C shows two examples and the others are shown in Sup. Fig. 7), showing that loss of recombination occurred independently and convergently for this region in the American and Eurasian lineages.

### 3. Dosage compensation of the Z-specific region

Many female-heterogametic species lack a chromosome-wide mechanism of dosage compensation, and investigating the few cases that have it may shed light on the difference between ZW and XY systems. Earlier work suggested that the Z-specific region of *A. franciscana* was compensated (Huylmans *et al*. 2019), but misidentification of genes in the sex-linked region (as the genome was fragmented) could have hidden differences between chromosomes. We repeated this analysis using RNA-seq data from thorax, head, and gonad of *A. sinica* (Huylmans et al. 2021). We first assembled a male transcriptome from all pooled male reads available for this species (to avoid hybrid assemblies of Z and W homologs, see Sup. Fig. 8 for a BUSCO assessment), mapped it to the genome assembly, and estimated expression for each sample. In somatic tissues, the female:male ratio is similar for the autosomes and each of the Z-chromosome regions (p>0.05, Fig. 2B and 2C), confirming that dosage compensation is active in this clade. A small shift towards male-biased expression can be observed for the S0 in gonads (Fig. 2D). Such differences in the gonad have been found even in animals with well-characterized chromosome-wide mechanisms of dosage compensation, such as Drosophila (Meiklejohn et al. 2011) and silkworm (Huylmans et al. 2017). While compensation mechanisms may be absent or less active in the gonad (Meiklejohn et al. 2011), differences could also result from the unusual regulation of the sex chromosomes in the germline (Argyridou and Parsch 2018), where they are often inactivated or downregulated (Vibranovski et al. 2009). Overall, our patterns generally support the presence of complete dosage compensation throughout the differentiated region in somatic tissues, and at least partial compensation in the gonad.

**Figure 2.**
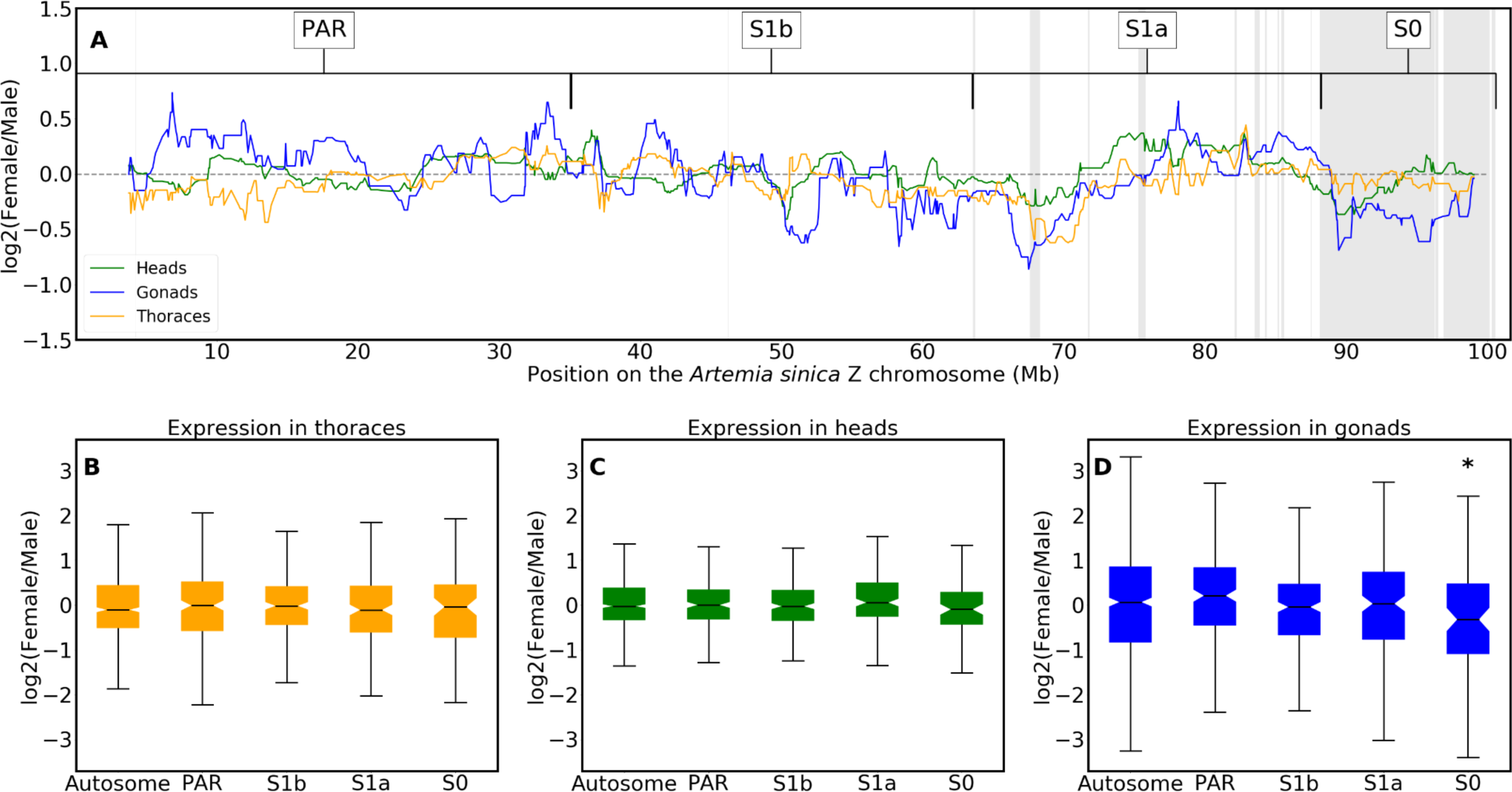
Dosage compensation of the Z-chromosome. Panel A: The logged ratio of female to male expression along the Z chromosome in heads, gonads and thoraces (computed as the rolling median in sliding windows of 30 consecutive genes). Gray areas represent the differentiated regions identified in the coverage analysis, and the putative strata are denoted above.The dashed horizontal black line is at zero. Panels B-D: The distribution of logged ratio of female to male expression for the autosomes and the different regions of the Z chromosome in thoraces (A), heads (B) and gonads (C).

### 4. The sex chomosomes of asexual females and the genetic origin of rare males

In order to characterize the ZW pair of asexual females, we first obtained a draft genome assembly of the closely related sexual species *A. sp. Kazakhstan* from illumina short reads, and estimated genomic coverage using two female and two male samples of this species. The genomic scaffolds were mapped to the *A.sinica* genome based on their gene content, and median coverages of male and female *A. sp. Kazakhstan* individuals were plotted along the *A. sinica* Z chromosome using a sliding window of 10 scaffolds (green and yellow lines in Fig. 3A). As expected, an approximately two-fold drop in female coverage was observed in a similar region to that found in *A. sinica* (marked by gray shading), whereas the male harbored high genomic coverage throughout the chromosome, consistent with the presence of the same pair of sex chromosomes in this lineage (a similar pattern was observed in *A. urmiana*, Sup. Fig. 9). We used the *A. sp. Kazakhstan* draft genome to map genomic reads derived from three closely related asexual females (one from the Lake Urmiana-derived population, and two from a population derived from Aibi Lake cysts). In every case, the patterns of coverage were very similar to those of the *A. sp. Kazakhstan* sexual female, confirming that asexual females carry the same pair of ZW chromosomes.

**Fig. 3:**
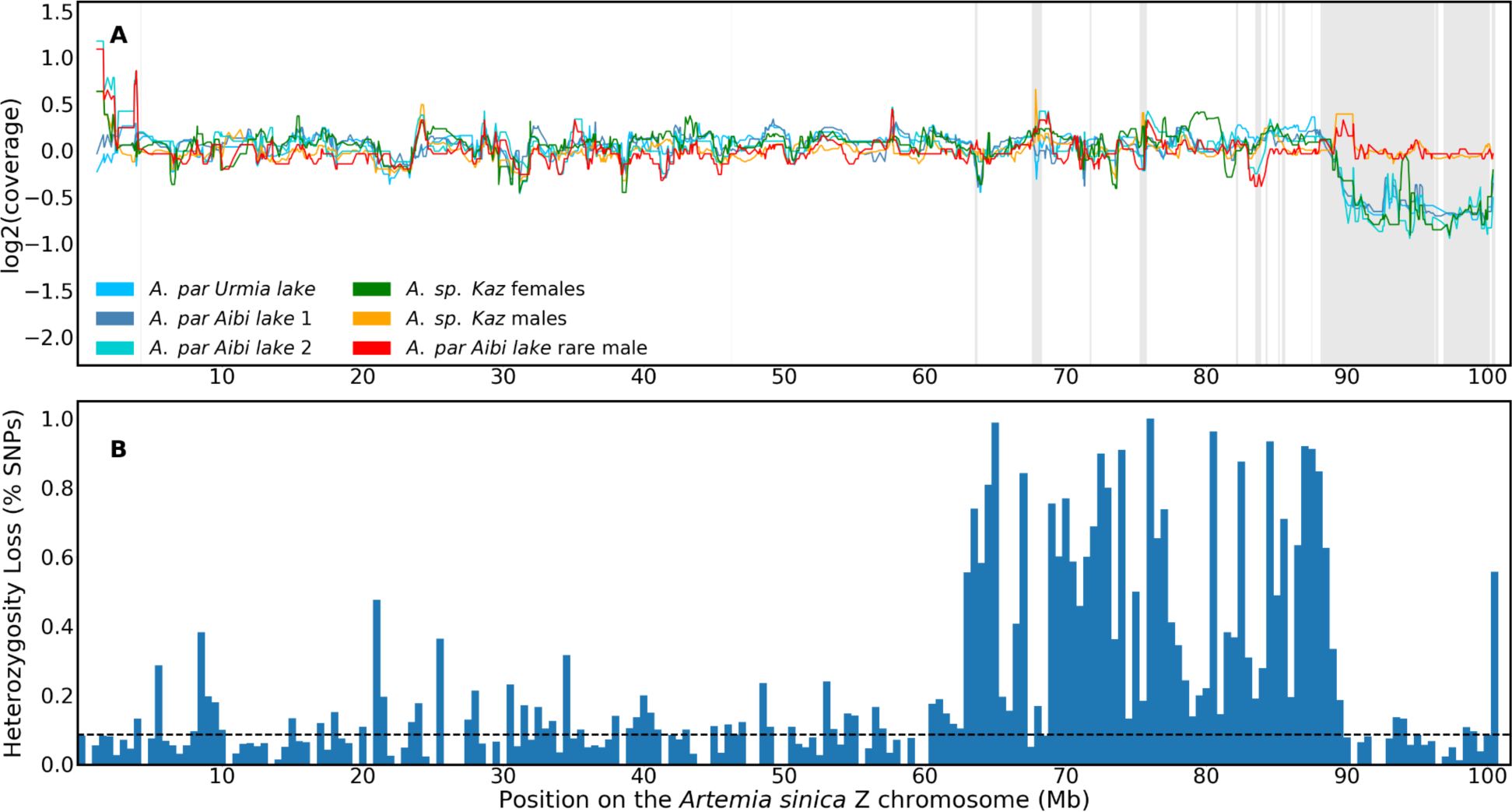
The sex chromosomes of sexual and asexual individuals. A. Coverage patterns in *A. sp. Kazakhstan* male and female samples, in three asexual females, and in a rare male derived from an asexual lineage from Aibi Lake. B. The fraction of SNPs that lost heterozygosity on the rare male Z chromosome relative to its asexual sister in bins of 500Kb. The dotted line represents the average loss of heterozygosity for autosomes.

Boyer *et al*. (2022) recently showed that *Artemia* rare males can be produced by ZW recombination events at variable locations near the sex-determining locus. We obtained a rare male from an *A. parthenogenetica* line from Aibi Lake (which we use in the next section to explore the transmission of asexuality). To test whether it arose through ZW recombination or other chromosomal changes, we first compared patterns of genomic coverage to those of females. No reduced coverage was observed along the Z-chromosome, arguing against the loss of a sex chromosome. We further called Single Nucleotide Polymorphisms (SNPs) in the rare male and in its sister (marked as Aibi female 2 in Fig 3A), and estimated the proportion of heterozygous SNPs present in the asexual female that were lost in the rare male. Loss of heterozygosity was detected throughout the distal half of the Z-chromosome (Fig. 3B and Sup. Fig. 10), confirming that a large part of the W was replaced by its Z homologous region. Finally, the rare male had high heterozygosity levels on other chromosomes (Sup. Fig. 11), arguing against the accidental occurrence of a version of automixis that eliminates variants across the genome (another hypothesis for the origin of rare males, Nougué *et al*. 2015). Taken together, these results support rare ZW recombination as the source of the Aibi Lake rare male (Nougué *et al*. 2015; Boyer *et al*. 2022).

### 5. The Z chromosome likely contributes to the transmission of asexuality

In order to find possible loci responsible for the spread of asexuality in brine shrimp, we crossed the rare male described in the previous section and a female from *A. sp. Kazakhstan* (Sup. Fig. 12). This produced 22 asexual females and 24 males in the F1; a single female died without producing offspring asexually. The presence of asexual females in the F1 shows that the locus controlling asexuality in this lineage works in a dominant manner, unlike what was first observed in Maccari *et al*. (2014), but consistent with the recent experiments of Boyer *et al*. (2021). The fact that almost all females produced offspring without mating further suggests that the locus was likely present on both copies of the genome of the original rare male. We then backcrossed 12 males from the F1, which should only carry one copy of the locus/loci controlling asexuality, with females from an *A. sp. Kazakhstan* inbred line (of these only 6 yielded progeny). The resulting F2 generation consisted of 84 (∼45%) males, 5 (∼3%) asexual females, and 96 (∼52%) females that did not produce asexually 133 days after the crosses were set up (44 individuals died before sexing was possible and are not included in the counts). We presume that most of these are sexual females for our analyses, but some could have reproduced asexually had the experiment been continued longer.

We produced whole-genome resequencing data for the 5 F2 asexual females and 10 F2 putatively sexual females. These were first pooled into an asexual pool and a putatively sexual pool, and we used Popoolation2 to compute FST between these two pools of females. While a few small peaks of FST are found on the autosomes (Fig 4a), the strongest signal comes from the distal end of the Z chromsome (Fig 4b). We further predicted that loci underlying asexuality should have been inherited from the original rare male by all the F2 asexual females, but not by (all) control females. To test this, we mapped all DNA samples individually to the *A. sp. Kazakhstan* genome. We also mapped the original rare male and its *A. parthenogenetica* sister, and two *A. sp. Kazakhstan* individuals, in order to select SNPs that were alternatively fixed between the two lineages. We used these informative SNPs to reestimate FST between F2 asexual and control females, and to infer which genomic regions were inherited from the rare male by each of the F2 individuals. Sup. Fig. 13 shows that we recover a region of high FST on the Z chromosome, and that all asexuals carry genetic material from the rare male in this region, as expected if it controls asexuality. In total, only 17 scaffolds show ancestry patterns consistent with an asexuality locus (i.e. they show evidence of *A. parthenogenetica* ancestry in all asexual females, but not in all control females). Eleven are on the Z chromosome (versus 1 expected, p=1.3e-20 with a Chi-square test), and correspond to the region of high FST, providing further support for a role of the Z chromosome in the transmission of asexuality. None of the other minor peaks of FST are in regions with ancestry patterns consistent with asexuality loci (Sup. Fig. 13), although chromosome 6 contains 3 such loci (versus 0.9 expected, p<0.01 with a chi-square test).

**Fig 4.**
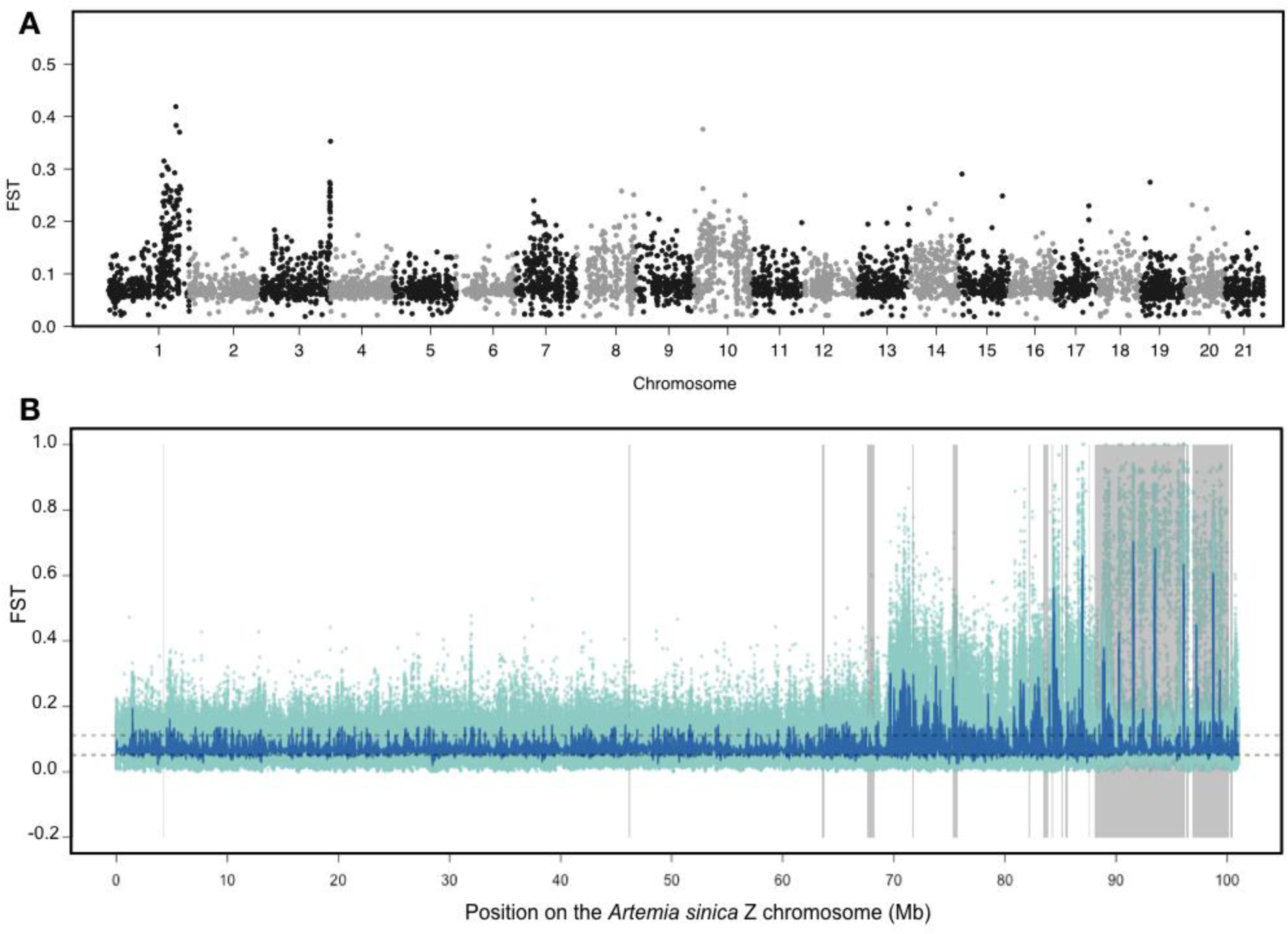
Elevated FST between sexual and asexual females localizes to the non-recombining region of the Z chromosome. A. Manhattan plot of FST estimated for 1Kb sliding windows between asexual and sexual females across the genome. B. FST across chromosome 1. FST is shown for individual SNPs in light green, and the blue line shows the rolling median for 101 SNPs.

## Discussion

Because of their unusual ecology (Gajardo and Beardmore 2012) and ease of maintenance in the lab, *Artemia* have for decades been used as model organisms for various purposes, including toxicity screening and ecological monitoring (Rajabi *et al*. 2015; Ntungwe N *et al*. 2020; Chan *et al*. 2021; Okumu *et al*. 2021), investigating molecular mechanisms of stress resistance (MacRae 2003), and even space research (Spooner *et al*. 1992). However, their potential as models for ZW chromosome evolution and comparative genomics in general was until recently hampered by a lack of genomic resources. The publication of two genomes for the American *A. franciscana* has already shed new light on how these charismatic organisms survive in their extreme environments (Jo *et al*. 2021a; De Vos *et al*. 2021), but no information on sex linkage was provided, and the lack of a close outgroup sequence (other than the distant Daphnia) made comparative analyses difficult. A draft genome was recently described for *A. sp. Kazakhstan* (Boyer *et al*. 2022), but there was limited power to assign scaffolds to the sex chromosomes or autosomes.

Here, we obtain the first chromosome-level assembly in the clade for the Eurasian brine shrimp *A. sinica*, and characterize in detail the differentiated and undifferentiated regions of the ZW pair. By combining these results with those of a preliminary analysis in *A. franciscana* (Huylmans *et al*. 2019), we confirmed the putative evolutionary model for the ZW pair, with an ancient well-differentiated region that stopped recombining in the ancestor of the two lineages, and more recent “strata” arising in each lineage independently. The independent loss of recombination in American and Eurasian species provides a unique opportunity to investigate convergent changes that occur in early sex-chromosome evolution. In agreement with previous findings in *A. franciscana*, *A. sinica* males and females have similar somatic expression patterns of Z-linked genes in the differentiated region which strongly supports the presence of a mechanism of dosage compensation in this group. Currently, no tractable lab model exists for the early evolution of ZW chromosomes, and Z-chromosome dosage compensation is only understood in detail for the silkworm (Walters and Hardcastle 2011; Kiuchi *et al*. 2014; Katsuma *et al*. 2019; Rosin *et al*. 2022), making this an outstanding new model clade for investigating these topics.

Finally, we obtained several putative W-genes in both species using a k-mer based analysis. Few of them mapped to the ancestral part of the W chromosome: only ∼20% of the Z-linked genes in this region have a W-homolog, suggesting that much of the ancestral gene content has been lost. All of the genes for which a W-homolog could be found in both *A. sinica* and *A. franciscana* mapped to younger strata of the ZW pair, and appear to have become W-linked independently in the two lineages. If the ancestral sex-determination mechanism is still shared by the two species, this may suggest that the primary signal for sex determination is a dosage-dependent gene on the Z rather than a dominant female-determining gene on the W. However, it is also possible that the sex-determining gene is only expressed early in development, and was missed by our analysis of adult tissues. Future studies of sex-specific expression throughout the life cycle and complete assemblies of the W chromosome of the two species will be necessary to shed light on sex determination in this group.

The proximity of *A. sinica* to the *A. sp. Kazakhstan* group, which contains both sexual and asexual populations, also allowed to characterize sex-linked sequences in this group. First, we found that the sex chromosome pair is shared by all populations. We further confirmed that rare males in this group can be produced through the replacement of the W-specific region with its Z-counterpart. Finally, our backcrossing experiment points to a role of the sex chromosome pair in the spread of asexuality through rare males.

It should be noted that this experiment has several drawbacks. First, it is difficult to phenotype females as sexual or asexual, as the timing at which asexuals produce their first brood can vary. Furthermore, hybrid incompatibilities may stop females from producing viable offspring even if they carry the alleles encoding asexuality. The fact that only ∼5% of females were asexual in the F2 suggests that either the trait is polygenic, and/or that we are mistakenly classifying asexuals as putative sexuals. Finally, we only obtained a small number of backcrossed asexual females, which limits the power to infer causal loci.

Despite these drawbacks, the Z chromosome showing the strongest signal of differentiation between asexual and control females is intriguing, and in line with results in the pea aphid, which carries the asexuality locus on the X chromosome. In species where asexuality is triggered by an endoparasite such as Wolbachia, the acquisition of asexuality is thought to be driven by the transmission advantage gained by the female-transmitted parasite (since asexual reproduction leads to an all-female progeny). It is possible that an asexuality gene found on a Z chromosome similarly benefits from a transmission advantage. If rare males always arise through the replacement of the W-specific region with its Z homolog region, a Z-linked asexuality locus will be homozygous and therefore transmitted to all daughters in the F1 (and to all sons). More detailed studies of transmission of asexuality in this group and others with ZW and XY sex chromosomes will in the future shed light on the relationship between sex determination and the rise and spread of asexual reproduction under various sex-determining mechanisms.

## Materials and methods

### Data availability statement

All genomic reads generated for this study are available at the NCBI short reads archive under Bioproject number XXX [*will be provided before publication*]. The pipelines used to analyze the data are at https://git.ist.ac.at/bvicoso/zsexasex2021, and important processed data files such as the new *A. sinica* genome assembly are provided in the Supplementary Materials. *[A permanent URL in the ISTA Data Repository will be provided before publication.]*

### Sampling and DNA extractions

Cysts from *A. sinica* (originally from Tanggu salterns, PR China), *A. sp. Kazakhstan* (originally from an unknown location in Kazakhstan), and two lineages of *A. parthenogenetica* (from Lake Aibi (PR China) and from Lake Urmia (Iran)) were obtained from the Instituto de Acuicultura de Torre de la Sal (C.S.I.C.) Artemia cyst collection in Spain, as described in Huylmans *et al*. (2021). Cysts were hatched under 30 g/L salinity and grown to adulthood under 60 g/L salinity. Some of these F0 individuals were used directly for DNA extractions with the Qiagen DNeasy Blood & Tissue kit. We also set up iso-female lines in *A. sinica* and *A. sp. Kazakhstan*, and subjected them to 6 generations of sib-sib mating to reduce the amount of heterozygosity. Male and female individuals from *A. sinica* and *A. sp. Kazakhstan* inbred iso-female lines were used individually for DNA extractions with Qiagen DNeasy Blood & Tissue kit. Furthermore, 20 males and 17 females of *A. sinica* (also from the inbred iso-female line) were pooled and high molecular weight DNA was extracted using the Qiagen Genomic-tip 20/G kit.

### Crossing design to identify the asexuality locus

We designed a backcross in order to investigate the loci controlling asexuality (Fig S12). An asexual female from Aibi Lake produced a rare male. We crossed this male with an inbred female from the closest related sexual species, *A. sp. Kazakhstan*. This produced asexual females and males in the F1 generation. We then backcrossed 12 males from the F1 to sexual females from the same inbred line of *A. sp. Kazakhstan*. Of these, 6 crosses produced offspring, yielding a total of 84 males, 5 asexual females, and 96 putatively sexual females (those that did not reproduce asexually for 133 days after the crosses were set up). The 5 asexual females and 10 control females were used individually for DNA extractions with the Qiagen DNeasy Blood & Tissue kit. The control females came from the same crosses (i.e. had the same F2 father and *A. sp. Kazakhstan* mothers) as the asexual females, but were otherwise selected randomly.

### DNA short and long read sequencing

PacBio libraries were prepared and sequenced at the Vienna Biocenter Sequencing facility for the male and female *A. sinica* high molecular weight DNA. All other DNA samples were used for Illumina paired-end sequencing. Libraries were prepared and sequenced at the Vienna Biocenter Sequencing Facility. Finally, 1 male was frozen and provided to the sequencing facility for Hi-C library preparation and Illumina sequencing on a NovaSeq machine. The final list of samples, as well as the parts of the analysis that they were used in, are listed in Sup. Table 3.

### Genome assemblies

The male PacBio reads were assembled using two different genome assemblers: Flye v.2.7.1, (Kolmogorov *et al*. 2019) and Miniasm [0.3-r179, minimap2 2.18-r1028-dirty was used for mapping and the consensus was generated using Racon v1.4.22](Li 2016; Vaser *et al*. 2017). The Flye assembly was polished using male *Artemia sinica* short genomic reads (trimmed with the Trimmomatic package, Bolger *et al*. 2014), and the Miniasm assembly was polished using the same male short reads using the wtpoa-cns tool from wtdbg2 v.2.5 (Ruan and Li 2020). The two assemblies were then merged using quickmerge v.0.3 (Chakraborty *et al*. 2016) with the Miniasm assembly as the query and the Flye assembly as the reference. The resulting assembly was purged using the purge_dups program v.1.2.5 (Guan *et al*. 2020)).

To scaffold the assembly into pseudo-chromosomes, the PCR duplicates were first removed from the Hi-C data using the clumpify.sh script from the bbmap package (Bushnell 2014), and the Hi-C reads were then mapped to the genome assembly using the Arima mapping pipeline with MAPQ 5 (Arima Genomics 2021) and then scaffolded using the YaHS tool (pre-release of version 1.1, Zhou 2022). The contact maps were visualized and manually edited on Juicebox v.1.11.08, Robinson *et al*. 2018) to generate the final chromosome-level assembly.

The female *Artemia sinica* genome was assembled from female PacBio reads using Flye (v.2.7.1), and it was not polished to avoid collapsing the Z and the W scaffolds. The *Artemia sp. Kazakhstan* genome was assembled from two male short read libraries with Megahit v1.1.4 (Li *et al*. 2015) and then scaffolded using SOAPdenovo-fusion (SOAPdenovo2 v.2.04, Luo *et al*. 2012).

BUSCO v.5.2.2, (Manni *et al*. 2021) was used to assess the completeness of the genomes generated in this study and the two previously published *Artemia franciscana* genomes in the genome mode with the arthropoda dataset (arthropoda_odb10).

### Estimation of genomic coverage

The short genomic reads were mapped to the genome using bowtie2 v.2.4.4 (Langmead and Salzberg 2012 p. 2). The uniquely mapped reads were then extracted from the output sam files using (grep -vw “XS:i”). SOAP.coverage v.2.7.7 (Luo *et al*. 2012) was then used to calculate the coverage for each library either using 10000 bp windows (*A. sinica*) or per scaffold (other species).

### Mapping of the *A. franciscana* and *A. sp. Kazakhstan* genomes to the new *A. sinica* assembly

We aligned the *A. sinica* published transcriptome (Huylmans *et al*. 2021) to both the *A. franciscana* and to the *A. sp Kazakhstan* genomic scaffolds using blat (Standalone BLAT v. 36x2, Kent 2002). For each transcript, we kept only the mapping location with the highest score in each genome (using the customized script 1-besthitblat.pl). When multiple transcripts overlapped by more than 20bps on the genome, only the transcript with the highest mapping score was kept (2-redremov_blat_v2.pl). We then used the location of the transcripts on the *Artemia sinica* genome to infer the location of the *A. franciscana* and *A. sp. Kazakhstan* scaffolds based on the transcripts they carried (AssignScaffoldLocation.pl). This script uses a majority rule to assign each scaffold to a chromosome, and then the mean location of genes on that scaffold to infer its final coordinate on the chromosome. All scripts are available on our Git page.

### FST between male and female populations

RNA-seq reads from 10 pooled *A. sinica* males and 10 pooled *A. sinica* females (from Huylmans et al, 2021), sampled from head, thorax and gonads, were pooled by sex and mapped separately to the male *A.sinica* reference genome using STAR (Dobin *et al*. 2013) with default parameters.

The resulting alignments with MAPQ score lower than 20 were filtered out and the remaining alignments were sorted using samtools view and sort functions (Li *et al*. 2009). Next, a pileup file of male and female alignments was produced using samtools-mpileup function. Finally, we used scripts from the Popoolation2 package (Kofler *et al*. 2011) to calculate FST. The mpileup file was reformatted with the Popoolation2 mpileup2sync.pl script, and the resulting synchronized file was used as an input for fst-sliding.pl script. FST between male and female populations was calculated for windows of 1000 nucleotides, using the fst-sliding.pl script with following options --suppress-noninformative --min-count 3 --min-coverage 10 --max-coverage 200 --min-covered-fraction 0.5 --window-size 1000 --step-size 1000 --pool-size 10.

### Identification of candidate W-genes with Kmer analysis

We used a k-mer based subtraction approach (Elkrewi et al., 2021) based on the BBMap package (Bushnell, 2014) on male and female genomic and RNA-seq data from *A. franciscana* and *A. sinica*. The pipeline was applied to each species separately. In *A. sinica,* two male and two female DNA libraries and two whole body RNA-seq replicates for each sex were used (Sup. Tables 3 and 4). In *A. franciscana*, the analysis was performed using one male and one female DNA libraries and pools of two RNA-seq replicates of heads and gonads for each sex, along with one whole body male and female RNA-seq libraries (SRR14598203 and SRR14598204).

First, the shared 31-mers between the female DNA and RNA libraries were identified, and then any k-mers matching male libraries were removed. Female RNA-seq reads containing these female-specific Kmers [with minimum kmer fraction of 0.6 (mkf=0.6)] were retrieved and assembled using Trinity (Grabherr *et al*. 2011), and the perl script from the Trinity package (get_longest_isoform_seq_per_trinity_gene.pl) was used to keep only the longest isoform. The male and female genomic libraries were mapped to the assembled transcripts using Bowtie2 (Langmead and Salzberg 2012 p. 2), and candidates with a sum of female perfect matches <=8 and a ratio of sum-of-females/sum-of-males <=2 were removed. The final set consisted of 402 transcripts in *A. franciscana* and 319 in *A. sinica*.

### Mapping of W candidates to the *A. sinica* genome

The *A. sinica* and *A. franciscana* W candidates were mapped to the *A. sinica* genome assembly with Parallel Blat (Wang and Kong 2019) with a translated query and database, and a minimum match score of 50. Only the mapping location with the strongest match score was considered for each transcript.

### Transcriptome assemblies and expression analysis

The *A. sinica* male transcriptome was assembled from two replicates of male whole body RNA-seq data (Huylmans *et al*. 2021) using Trinity (Grabherr *et al*. 2011) in two different modes: denovo and genome-guided. The two assemblies were concatenated and then filtered using the tr2aacds.pl script from EvidentialGene (Gilbert 2019). For the expression analysis, only the first isoform was kept for each gene, and only transcripts longer than 500bp were used in the analysis. The RNA-seq reads from the *A. sinica* heads, gonads, and thoraces of males and females (Huylmans *et al*. 2021) were mapped to the curated transcriptome and TPM values were obtained using Kallisto v.0.46.2, (Bray *et al*. 2016). Normalization was done using NormalyzerDE (Willforss *et al*. 2019).

Two different *A. franciscana* de novo transcriptome assemblies were made using Trinity. The first using pooled RNA-seq reads from male heads and testes (two replicates each, Huylmans *et al*. 2019), and the second using the published whole-body male RNA-seq library (SRR14598203, Jo *et al*. 2021b). The two assemblies were concatenated and then filtered using the tr2aacds.pl script from EvidentialGene.

### Phylogenetic Trees

The W candidates of *A. sinica* and *A. franciscana* were mapped reciprocally to each other using pblat (BLAT with parallel supports v. 36x2 with default parameters, Wang and Kong 2019), and reciprocal best hits were considered shared candidates. The W candidates of the two species were further mapped to their respective uncollapsed male transcriptome assemblies (see previous section) with pblat (Wang and Kong 2019) with a translated query and database, and a minimum match score of 50. The transcripts with the highest mapping score to the W candidates were used as the putative Z homologs in their respective species.

The *Branchinecta lindahli* transcriptome (Schwentner *et al*. 2018) was downloaded from the Crustacean Phylogeny dataset on Harvard Dataverse (https://doi.org/10.7910/DVN/SM7DIU). *B. lindahli* homologs of shared W-candidates were obtained by mapping the putative Z homologs of both species to the *B. lindahli* transcriptome using pblat (-minScore=50 -t=dnax -q=dnax) and retrieving the transcript with highest alignments score (using the customized script 2-besthitblat.pl). A transcript was considered a homolog if it mapped to at least one of the putative Z homologs of the two species, and when the two Z homologs mapped to different outgroup sequences, both outgroup sequences were retrieved and used to make two different alignments.

The shared W candidates of *A. sinica* and *A. franciscana*, their Z homologs, and the outgroup sequences were aligned using MAFFT (v7.487, with the options “mafft -- adjustdirection INPUT > OUTPUT”, Katoh *et al*. 2002). The resulting alignments were fed to phylogeny.fr (Dereeper *et al*. 2008), where the alignment was curated using GBLOCKS (Talavera and Castresana 2007), and the phylogenetic tree was constructed using PhyML (Guindon *et al*. 2010). Trees were made only for sequences where the number of overlapping positions after gblocks was longer than 200bp (Sup. Fig. 7). In the four instances where the curated alignment length with the outgroup was shorter than 200bp, we tried aligning the sequences without the outgroup. For the two cases where the resulting alignment length was longer than or equal 200bp, unrooted trees were made (Sup. Fig. 7). The trees were then downloaded in the Newick format and visualized using itol.embl.de (Letunic and Bork 2019).

### Heterozygosity analysis in asexual female and rare male

Illumina genomic sequencing was performed on a rare male and its asexual sister (both derived from an Aibi Lake *A. parthenogenetica* lineage), yielding around 115 million paired-end reads with a length of 125 nucleotides for each sample. The reads were quality- and adapter-trimmed with Trimmomatic-0.36 (Bolger *et al*. 2014), and mapped to the draft *Artemia sp. Kazakhstan* genome assembly using STAR v.2.6.0c (Dobin *et al*. 2013) with default settings.

We indexed the reference *A. sp. Kazakhstan* genome using SAMtools v.1.10 (Li *et al*. 2009), called the SNPs from BAM alignments with BCFtools v.1.10.2 (Li *et al*. 2009), then removed indels, filtered for quality of reads over 30 and coverage over 5 and below 100 with VCFtools v.0.1.15 (Danecek *et al*. 2011), and removed multiallelic sites with BCFtools.

We calculated the fraction of SNPs that lost heterozygosity in the rare male DNA in comparison with the asexual sister DNA. It was calculated and visualized in 500kb bins for each chromosome.

### Analysis of backcross between the Aibi Lake rare male and *A. sp. Kazakhstan* females

We sequenced 5 asexual females and 10 putatively sexual females from the F2 generation. This resulted in an average of 101 million reads per asexual female and 50 million reads per putatively sexual female. We removed adaptors and trimmed reads using Trimmomatic v0.39 (Bolger *et al*. 2014). We then aligned the resulting paired-end reads to the genome using Bowtie2 v2.4.4 (Langmead and Salzberg 2012). SAM files were converted to BAM files and sorted in Samtools v.1.13 (Li *et al*. 2009).

For our pooled analyses, we merged BAM files into a pooled asexual BAM file and a pooled putatively-sexual BAM, and created a mpileup file in Samtools v.1.13. We then used Popoolation2 (Kofler *et al*. 2011) to call FST for both individual SNPs and in 1kb windows. We used FST computed for 1kb windows to visualize FST across the genome in a Manhattan plot in the R package qqman (Turner 2018). We computed rolling medians in sliding windows of 101 consecutive SNPs on each linkage group using the rollmedian function from the package zoo (Zeileis and Grothendieck 2005) in R v.4.0.3. To identify regions of elevated FST on individual chromosomes, we computed 95% confidence intervals by sampling rolling medians of 101 consecutive SNPs across the genome 1000 times.

For our individual-based analyses, we first used SEQTK v1.2 (https://github.com/lh3/seqtk) to randomly select a subset of reads from each asexual sample to match the highest coverage found in an F2 control female (to avoid biases caused by the much larger number of reads obtained for the F2 asexuals than for the controls). We then mapped reads from all F2 individuals to the *A. sp. Kazakhstan* genome using BWA mem v0.7.17 (Li and Durbin 2009) with default parameters. DNA reads from the rare male and its *A. parthenogenetica* sister, and from two *A. sp. Kazakhstan* individuals, were also subsetted and mapped. The resulting BAM alignments were sorted with samtools v1.14 (Li *et al*. 2009), and used to call SNPs with the mpileup function of BCFtools v1.14 (Li 2011). The VCF file was filtered with VCFtools v0.1.17 (Danecek *et al*. 2011) for minimum and maximum depth (4 and 50), minimum quality score (30) and minimum minor allele frequency (0.1). Only SNPs for which the two *A. sp. Kazakhstan* had a genotype of 0/0, and the two *A. parthenogenetica* individuals 1/1, were kept for further analyses. We computed FST between the F2 asexual and control females using the function -- weir-fst-pop in VCFtools for 10kb windows. We then inferred ancestry of each genomic scaffold in every sample (i.e. whether they were homozygous for the *A. sp. Kazakhstan* haplotype, or carried a copy of the *A. parthenogenetica* haplotype as well) using the customized script Chromopaint.pl (available on our git page). The *A. sp. Kazakhstan* genomic scaffolds were assigned to a location on the *A. sinica* genome as before. Scaffolds with more than 10 informative SNPs, and >80% 0/1 or 1/1 SNPs were considered to be heterozygous for the *A. sp. Kazakhstan* and *A. parthenogenetica* haplotypes, whereas scaffolds with >80% 0/0 were considered to have only *A. sp. Kazakhstan* ancestry (only 5 to 9% of scaffolds fell in between and could not be classified in each individual).

## Acknowledgements

We thank the Vicoso group for comments on the manuscript, and the ISTA Scientific computing team and the Vienna Biocenter Sequencing facility for technical support. This work was supported by the European Research Council under the European Union’s Horizon 2020 research and innovation program (grant agreement no. 715257).

